# Exploring white matter dynamics and morphology through interactive numerical phantoms: The White Matter Generator

**DOI:** 10.1101/2023.12.08.570748

**Authors:** S. Winther, O. Peulicke, M. Andersson, H. M. Kjer, J. A. Bærentzen, T. B. Dyrby

## Abstract

Brain white matter is a dynamic environment that continuously adapts and reorganizes in response to stimuli and pathological changes. Glial cells, especially, play a key role in tissue repair, inflammation modulation, and neural recovery. The movements of glial cells and changes in their concentrations can influence the surrounding axon morphology. We introduce the White Matter Generator (WMG) tool to enable the study of how axon morphology is influenced through such dynamical processes, and how this, in turn, influences the diffusion-weighted MRI signal. This is made possible by allowing interactive changes to the configuration of the phantom generation throughout the optimisation process. The phantoms can consist of axons, myelinated axons, and cell clusters, separated by extra-cellular space. Due to morphological flexibility and computational advantages during the optimisation, the tool uses ellipsoids as building blocks for all structures; chains of ellipsoids for axons, and individual ellipsoids for cell clusters. After optimisation, the ellipsoid representation can be converted to a mesh representation which can be employed in Monte-Carlo diffusion simulations. This offers an effective method for evaluating tissue microstructure models for diffusion-weighted MRI in controlled realistic white matter environments. Hence, the WMG offers valuable insights into white matter’s adaptive nature and implications for diffusion-weighted MRI microstructure models, and thereby holds the potential to advance clinical diagnosis, treatment, and rehabilitation strategies for various neurological disorders and injuries.

## 1 Introduction

The neuronal network of the brain is a dynamic structure, constantly adapting to internal and external stimuli. Within the white matter, the local environment of neurons is found to modulate their axon morphology [1]. Especially glial cells play a significant role in this modulation. While the glial cells do not directly participate in neuronal signalling, they carry out crucial supporting tasks: myelination (oligodendrocytes), maintaining an appropriate chemical environment for neuronal signalling (astrocytes), and removing cellular debris from sites of injury or normal cell turnover (microglial) [2]. To carry out these tasks, the glial cells move through the tissue. Such dynamic behaviour has been observed with microscopy [3][4][5]. Since the morphology of glial cells generally does not change, primarily the morphology of the axons adapts [1].

Moreover, there is a relation between the morphology and function of the individual neurons. Morphological factors that affect the axonal conduction velocity include the axon diameter [6][7], the myelin sheath thickness [8], and the length and spacing of nodes of Ranvier [9]. Quantifying the morphology of axons and brain white matter in general is thus of great interest for enhancing our understanding of various brain processes, and can provide a potential biomarker of disease.

The most promising method for assessing axon morphology non-invasively, is diffusion-weighted MRI (dMRI). dMRI reflects the morphological properties of the underlying tissue by measuring diffusion properties of water molecules within the tissue [10][11][12][13]. To advance the method even further, we need access to extensive ground truth-informed experimentation for scan sequence optimization and biophysical model validation. This is unfeasible to obtain with either preclinical or clinical MRI due to the limited spatial resolution.

It is therefore of great interest to supplement dMRI research with numerical simulations, because these come with a ground truth. dMRI can with advantage be simulated based on Monte Carlo (MC) diffusion simulations [14][15]. The level of realism in simulations is crucial for the specificity and applicability obtainable by the models developed based on them [16][17][1][18].

Mimicking the highly complex tissue of white matter is a great challenge. White matter has commonly been represented as idealized straight, coaxial, infinitely long, and non-touching cylinders [13]. However, from recent advances in 3D histology, it has been validated that the white matter has a much more complex configuration [5][19][20][1]. While individual axons are found to express non-circular cross-sections, longitudinal diameter variations, and tortuosity, axons on the ensemble scale are found to express orientation dispersion and crossing fibres. Furthermore, recent studies show that such characteristics have a crucial impact on the diffusion signal and modelling [16][17][1][18][21], and should therefore be incorporated into our simulations to improve the realism.

Two primary approaches have been taken to generate realistic numerical white matter phantoms: segmentation (both semiautomatic and automatic) from 3D histology of tissue [19][22][20][1], and numerical synthesis [23][24][25][26][27][28][29][30]. Segmentation of white matter tissue provides a very high degree of realism. However, it can be very resource-consuming with respect to time for manual editing and the sacrifice of animal lives. Furthermore, this approach provides little flexibility in the morphological variation of the tissue and only provides static snapshots thereof. In contrast, numerical synthesis allows for much lower resource consumption and allows for a high degree of morphological flexibility. However, the outputs are based on assumptions of what the anatomy looks like [31].

Various tools have been developed for numerical synthesis of white matter phantoms with complex morphological properties. Recent tools include MEDUSA [28], ConFiG [29], and CACTUS [30]. MEDUSA [28] enables the representation of the most diverse tissue elements by allowing multiple compartments including axons, astrocytes, oligodendrocytes, nodes of Ranvier, and myelinated axons. All structures are represented as spheres. While the sphere representation allows for high representational power of the longitudinal axon morphology characteristics, it does not allow for eccentric cross-sections documented by histology [19][1][20]. Meanwhile, both ConFiG [29] and CACTUS [30] allow only for the inclusion of myelinated axons. However, due to a higher refinement of the axonal crosssections, these possess higher realism compared with the otherwise circular cross-sections. CACTUS [30] stands out for its superior computational efficiency, which in turn enables the creation of larger and more dense phantoms. However, none of these tools considers the dynamic aspects of white matter tissue which has a crucial influence on axon morphology [1].

In this work, we present the White Matter Generator (WMG); a new tool for generating interactive numerical phantoms for Monte Carlo dMRI simulations. The WMG tool enables the mimicking of brain white matter dynamics; inspired by high-resolution 3D synchrotron X-ray nanoholotomography (XNH) imaging in our previous work [1][32][33]. The phantoms can consist of multiple compartments: axons, myelinated axons (here defined as fibres), and cells, separated by extra-cellular space. The key assumption for the synthesis is that axon morphology is modulated by the local environment, as observed with XNH imaging in our previous work [1][32]. Due to morphological flexibility and computational advantages during the optimisation, the tool uses ellipsoids as building blocks for all structures during the synthesis; chains of ellipsoids for fibres, and individual ellipsoids for cells. Thereby, fibres can obtain non-circular cross-sections, longitudinal diameter variations, and tortuosity as observed in 3D histology. To mimic a dynamic tissue environment, the tool allows for interactive adaptation by changing parameters and tissue composition during the optimisation process. The output is in the format of PLY surface meshes, which makes them directly compatible with existing Monte Carlo diffusion simulators such as the widely used MC-DC Simulator [15] and Camino [14]. We demonstrate examples of various tissue configurations: varying degrees of fibre dispersion, types of bundle crossings, demyelination, inclusion of static cells, and cell dynamics. The biological relevance of the outputted axons is evaluated based on a set of morphological metrics and compared with those observed with XNH imaging of monkey brain white matter in our previous work [1][32].

## 2 Methods

Phantoms can consist of myelinated axons, axons, and cells, separated by extra-cellular space. To obtain morphological flexibility and computational advantages during the optimisation, the tool uses ellipsoids as the building blocks for all structures during the synthesis; chains of ellipsoids for fibres, and individual ellipsoids for cells.

The optimisation is based on a force-biased packing algorithm, first introduced by [34], where the phantoms are obtained as an equilibrium between *repulsion forces* (for avoiding overlap between individual axons and cells) and *recover forces* (for ensuring the structure of the individual axons) between the ellipsoidal building blocks.

Different brain regions and types of cell dynamics can be mimicked by adapting the configuration w.r.t. fibre diameter distributions, fibre volume fractions (FVF), cell volume fractions (CVF), global dispersion (*ϵ*), bundle crossings, and interactive changes. Changes to the configuration can be made interactively and at any time during the optimisation.

An optimisation starts with one configuration file (config-file) and ends with another updated config-file. Thereby, the updated config file can be easily adapted and given as input for another round of optimisation according to these adaptations. A conceptual flow chart is seen in Fig. 1, and each compartment is described in more detail later in this section.

**Figure 1:**
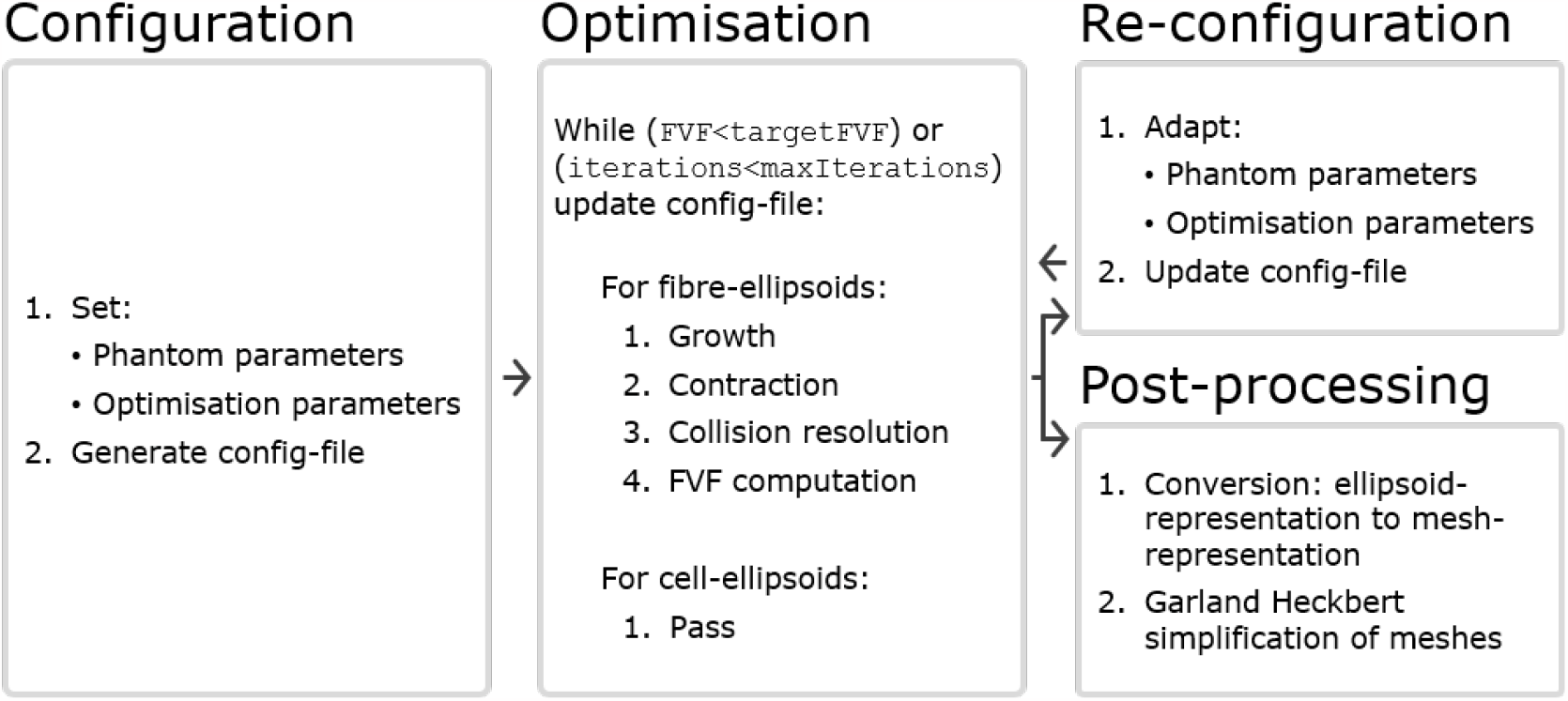
The WMG tool can be divided into four compartments. **Configuration:** A config-file is generated based on a set of phantom parameters and optimisation parameters. Within the config-file, all structures are represented by ellipsoids. **Optimisation:** The optimisation runs based on the config-file, and is carried out by iterating over the axonal ellipsoids. Cellular ellipsoids remain static. Maintaining consistency between the input and output format makes interactive adaptation of the configuration convenient. **Reconfiguration:** After a round of optimisation, the config-file can be adapted by changing phantom parameters (e.g. CVF and cell positions), and/or optimisation parameters (e.g. `ellipsoidDensity`, `growSpeed` and `maxIterations`). It can then be used as input for another round of optimisation. **Post-processing:** After a round of optimisation, the config-file can be post-processed to obtain a mesh-representation from the ellipsoid-representation. Furthermore, it is often beneficial to perform a Garland Heckbert simplification of these meshes to remove redundant vertices to reduce the computational load of e.g. Monte Carlo diffusion simulations.

**Figure 2:**
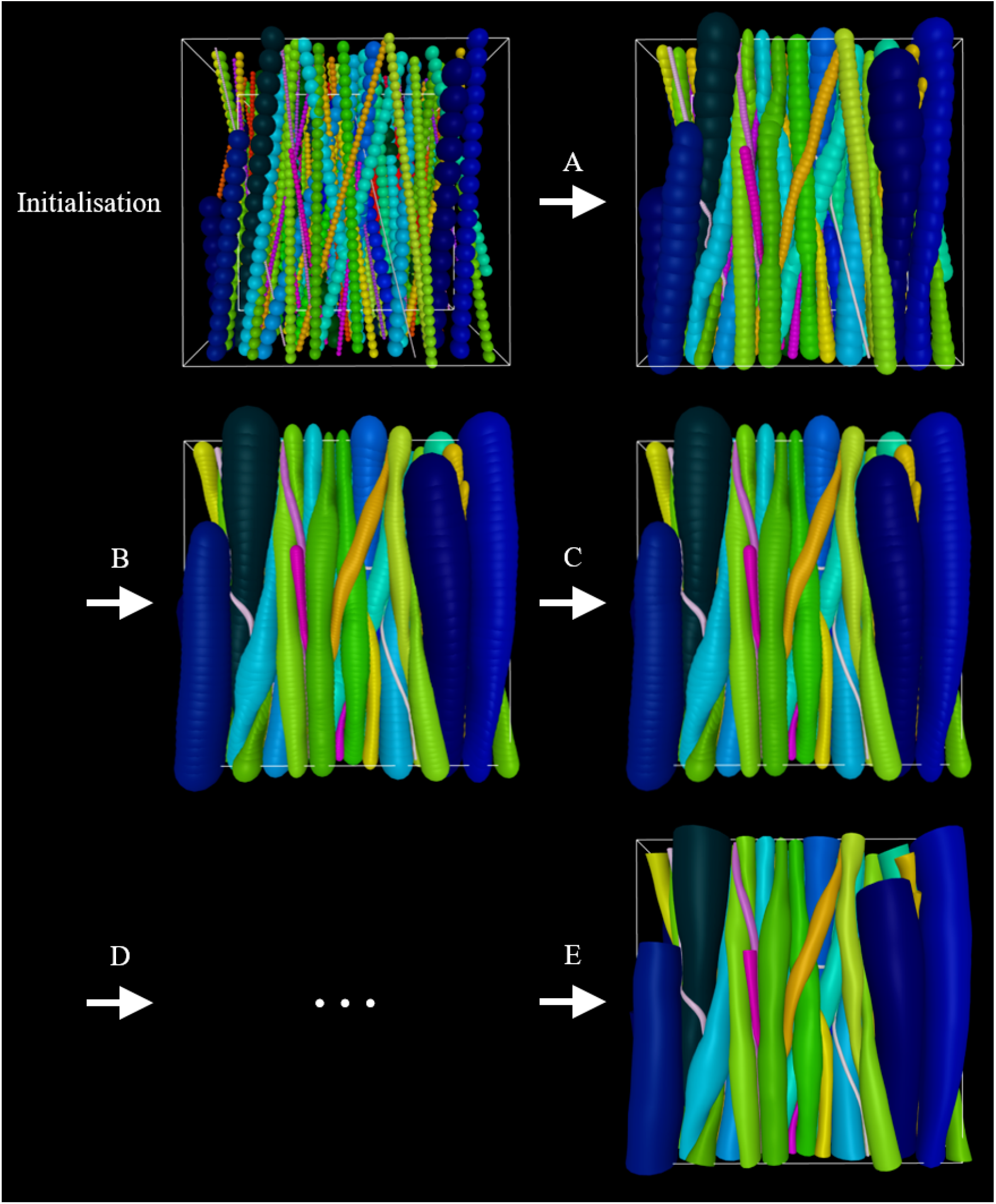
Example of an optimisation scheme. At **initialisation**, the phantom consists of an outer-voxel (for spatial restriction), an inner voxel (for the computing optimisation metric FVF), straight fibres, and cells. Initially, smaller diameters are assigned to the fibres to allow for more flexibility as they grow into their converged morphology. **A)** Inter-fibre ellipsoids grow step-wise towards their maximum allowed diameter while adapting to their surroundings as a consequence of collisions with these. **B)** The fibre-ellipsoid density is increased. **C)** Growth is repeated. **D)** Steps B) and D) or variations thereof can be repeated. **E)** Once optimised to the user’s liking, the ellipsoid-representation can be converted to a mesh-representation and then applied in e.g. Monte Carlo diffusion simulations.

The WMG tool is written in TypeScript, whereas supporting functions for configuration and post-processing are written in Python 3. A simplified version of the WMG tool is available as a graphical web interface at https://white-matter-generator.github.io/. This version allows one to get a more intuitive feeling of the tool and its parameters while testing out different configurations. The full version is available as a command line interface which can be installed by following the instructions at https://white-matter-generator.github.io/help/cli. This version enables automation and large-scale production, and this is the version we will be focusing on here. The supporting functions for configuration and post-processing will be made available through GitHub.

### 2.1 Configuration

There are two types of parameters: *phantom parameters*, which specify the content and limits of the phantom, and *optimisation parameters*, which specify the optimisation criteria of the phantom. As long as the config-file follows the correct formatting, the user can configure the phantom in any way they like. Below is a guide to the options provided with the WMG.

#### The phantom parameters are

- Voxel:

–Outer (larger) voxel dimensions: Defines the spatial restriction of phantom content as a cuboid.

–Inner (smaller) voxel dimensions: Defines the cuboid volume wherein the FVF is optimised. This voxel is necessary because the discontinuation at the outer voxel boundary will lead to biased morphology at that boundary.

- Fibres:

–Initial base point (**b**) and direction (**r**): Fibres are initialised as chains of ellipsoids arranged in straight lines defined with a direction and one fixed base point. The fixed points are uniformly distributed along the axis of primary fibre-orientation (i.e. the axis perpendicular to the idealised 2D plane).

–Global fibre dispersion (*ϵ*): Fibre directions are uniformly sampled on the surface of a spherical cone, where the height of the cone cap is scaled by *ϵ. ϵ* = 0.0 allows disperson of 0 deg and *ϵ* = 1.0 allows dispersion of 90 deg.

–Crossing bundles: Can be configured either as adjacent sheets [33] or interwoven. Specified by the number of bundles, their primary direction and the fraction of fibres they contain.

–Fibre-ellipsoid separation (`mapFromMaxDiameterToEllipsoidSeparation`): The ellipsoid separation within each fibre is defined from a user-specified mapping function which takes the maximum allowed fibre-diameter as input.

–Allowed diameter ranges for the individual fibres (`mapFromMaxDiameterToMinDiameter`): To obtain high packing the ellipsoids within each fibre can be allowed to vary in diameter. The range is defined by a user-specified mapping function which takes the maximum allowed fibre-diameter as input. We base the allowed range on an idealised packing of straight cylinders (corresponding to the packing of discs in a 2D plane). This packing is obtained from the MC-DC Simulator [15]. We initialise all fibres with a diameter which is significantly lower than the idealised case in order to achieve the more complex packing.

–g-ratio: When the ellipsoid-representation is converted to the mesh-representation the fibre is compartmentalised into axon and myelin based on this g-ratio. I.e. the myelin diameter matches that of the fibre, while the axon diameter is scaled in relation to the fibre’s diameter through the g-ratio.

–Contract speed (`contractSpeed`): How much the fibres contract per step, i.e. how stiff the axons are. This number should be ≥ 0.

–Deformation factor (`mapFromDiameterToDeformationFactor`): How much the fibre-ellipsoids should deform as opposed to change position when a collision occurs. The deformation is a number between 0 and 1, and is defined by a user-specified mapping function which takes the current fibre-diameter as input. A deformation factor of 0 means that the ellipsoid cannot be deformed at all and hence remains a perfect sphere. A deformation factor of 1 means that the ellipsoid will deform as much as possible rather than change the position of its centre.

- Cells:

–Target CVF (`targetCVF`): Cells can be added based on ellipsoidal dimensions randomly sampled from normal distributions. One cell is sampled at a time. Cells are added until the `targetCVF` is met.

–Position: Defines the center of the cell.

–Shape: Specifies the 3x3 transformation matrix used when going from a unit sphere to an ellipsoid. This ellipsoid will be the shape of the cell.

- Output format:

–Radial resolution of outputted meshes: Determines how detailed the output mesh will be.

–Extend fibres around the voxel: Defines whether or not to extend the fibres around the voxel to create mirrored intra-axonal compartments.

#### The optimisation parameters are

- Target FVF (`targetFVF`): The targeted FVF.
- Grow speed (`growSpeed`): How much the fibres grow per optimisation step. 0 means no growth, and 1 means that the axon will grow to 100% of its target size in 1 step.
- Maximum number of iterations (`maxIterations`): Even if the `targetFVF` is not reached at this point, the optimisation will terminate here.
- Output interval (`outputInterval`): Interval with which an output with the current configuration is provided.

It is beneficial to apply optimisation schemes that start out with higher `growSpeed` (bigger steps) and lower intra-fibre ellipsoid density (less computationally demanding), and go towards lower `growSpeed` (smaller steps) and higher intra-fibre ellipsoid resolution (more computationally demanding). Furthermore, it is crucial to choose realistic parameters to obtain a successful phantom optimisation.

### 2.2 Optimisation

#### 2.2.1 Initialisation of voxels, fibres and cells

Each fibre is initialised with a base point, **b**, and a direction, **r**. A ray is traced along **r** and − **r** until the voxel boundary is hit. The two points hit by the ray define the endpoints of the fibre. The straight line connecting the two endpoints is then filled with ellipsoids according to the specified ellipsoid density and minimum fibre diameter. Each cell is represented by a single ellipsoid. They cannot move or deform, but otherwise act in the same way as fibres. Each ellipsoid has two properties: a position vector and a shape matrix. With no deformation, the shape of an ellipsoid is a unit sphere *S*^2^ = {**q** ∈ ℝ ^3^|∥ **q** ∥= 1} . The deformation of the sphere is represented by a 3 *×* 3 matrix, **S**, such that the surface of the resulting ellipsoid is *E* = {**Sq** + **p**|∥ **q**∥ = 1} where **p** is the position.

Initially, the matrix **S** = *μ***I** where *μ* is given by the allowed minimum radius for the given fibre. For increased flexibility, *μ* should be very small relative to the voxel size.

Hence, each ellipsoid, *j*, on each fibre, *i*, is represented by a position vector, **p**_*i,j*_, and a shape matrix, **S**_*i,j*_, and the surface of the ellipsoid is thus defined by

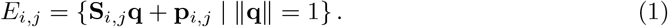

Ellipsoids are deformed by applying a deformation matrix **D** = *s***rr**^*T*^ + **I** . Multiplying (the points on) a surface by **D** thereby results in a scaling of 1 + *s* along **r** where *s* ∈ ℝ and **r** ∈ ℝ^3^, ∥**r**∥ = 1. This can be seen by taking an arbitrary point **p** and transforming it using **D**.

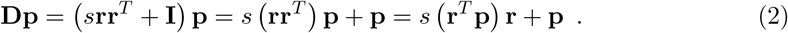

After *n* deformations of an ellipsoid the shape matrix will have the form **S** = **D**_*n*_…**D**_2_**D**_1_.

#### 2.2.2 Growth, contraction and redistribution for fibre-ellipsoids

The growth step considers each ellipsoid *j* of each fibre *i*. The shape matrix of the updated ellipsoid **S**_*new*_ is set to a weighted average between the current shape **S**_*old*_ and the target shape (i.e. a sphere having the maximum radius *r*_*max*_) by

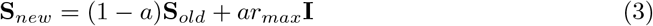

where *a*∈ [0, 1] is the `growSpeed`.

A contraction step adjusts the curve on which the ellipsoids lie. The force along the fibre is dependent on the contraction coefficient *κ*. A value of *κ* = 0 will do nothing, and a value of *κ* = 1 will move the position **p**_*i,j*_ of the ellipsoid all the way over to 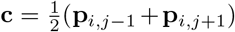 making this segment of the fibre a straight line. For any value of *κ* ∈ [0, 1] the position is updated to (1 − *κ*)**p**_*i,j*_ + *κ***c**.

A redistribution step is then performed to prevent ellipsoids from clumping together and thereby creating gaps in the ellipsoid chain. This is done by updating the position of each ellipsoid in a chain such that they are evenly distributed along the trajectory of the fibre. Furthermore, the endpoints of the fibres are moved to the nearest side whenever they are inside the voxel to make sure that the fibre spans the entire voxel.

#### 2.2.3 Collisions: fibre-voxel, fibre-fibre, and fibre-cell

We then deal with collisions. Fibre-voxel collisions (outer voxel) are checked for each ellipsoid independently. Whenever an ellipsoid’s center is outside of the voxel it is projected back onto the boundary. Fibre-fibre collisions are checked by computing the potential overlap between each ellipsoid of a fibre and all other ellipsoids of all other fibres. Fibre-cell collisions are checked by likewise computing the potential overlap between each ellipsoid of a fibre and all cells.

To compute overlaps between ellipsoids (see Fig. 3), we utilise that finding the surface point of an ellipsoid furthest away along a vector **r** is the same as finding the surface point which has the normal **r**. When deforming an ellipsoid using the matrix **D**, the normals are transformed by **D**^−1^. Since after *n* deformations **D**_*i*_, *i* ∈*{*1, …, *n}* the shape has the form **S** = **D**_*n*_…**D**_2_**D**_1_, the corresponding transformation matrix **N** for the normals is

**Figure 3:**
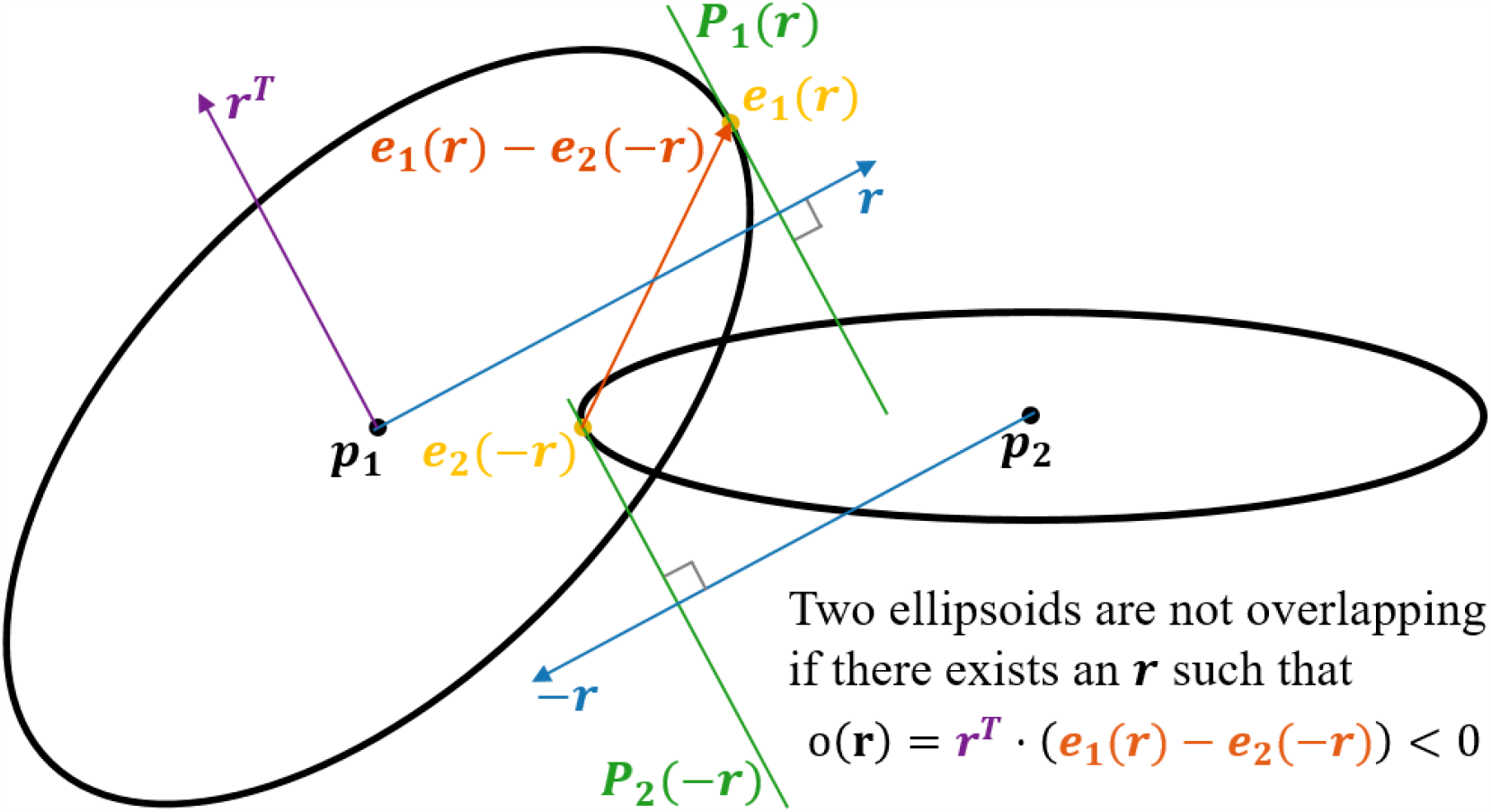
2D visualisation of the concept applied for checking for potential overlap of two ellipsoids. In the example, we have two ellipsoids centered at **p**_**1**_ and **p**_**2**_, respectively. The objective is to check if there exists a set of hyperplanes, **P**_**1**_(**r**) and **P**_**2**_(−**r**), which separates the ellipsoids. This is done by finding the surface point **e**_**1**_(**r**) for which the normal is parallel with *r*, and the surface point **e**_**2**_(−**r**) for which the normal is parallel with −**r**. Now, if the dot product between the vector **r**^*T*^ (lying in the plane **P**_**1**_(**r**) and the vector **e**_**1**_(**r**) − **e**_**2**_(−**r**) is less than 0, the planes **P**_**1**_(**r**) and **P**_**2**_(−**r**) must separate the two ellipsoids. I.e., if there exists an **r** such that **r**^*T*^ *·* (**e**_**1**_(**r**) − **e**_**2**_(−**r**)) *<* 0, the two ellipsoids are not overlapping.

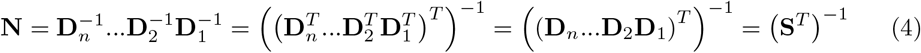

since each **D**_*i*_ is symmetric. When going from a normal on an ellipsoid to the corresponding normal of the original sphere, the normal should be transformed by the inverse, i.e. **N**^−1^ = **S**^*T*^ . After this transformation, the resulting vector can be normalized. Since we now have a unit normal on a unit sphere, the position on that sphere is given by the same vector. To get the corresponding position on the ellipsoid, one can simply multiply by **S**. Hence the point with the normal **r** on the ellipsoid with shape **S** and center at the origin is 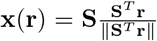 . Given the ellipsoid *j* of fibre *i* and a direction **r** the extremal point is thus **e**_*i,j*_(**r**) = **p**_*i,j*_ + **x**_*i,j*_(**r**). This extremum is used to compute the overlap between two ellipsoids.

Define the overlap of two ellipsoids *j*_1_ and *j*_2_ belonging to the fibres *i*_1_ and *i*_2_ respectively along an axis **r** as 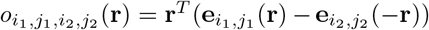. If one can find a *separating* axis [35], **r**, such that 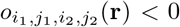, the ellipsoids don’t overlap. If 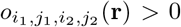 for all values of **r**, the ellipsoids overlap, since they are convex. To check whether two ellipsoids overlap, one can find the minimum overlap and check if it is positive or negative. I.e. the ellipsoids overlap if and only if 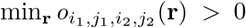. The minimum is computed numerically starting with the initial guess 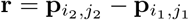 .

When the axis **r** providing the least overlap 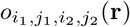 is found, it is checked if 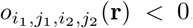. If so, nothing should be done. In the case where 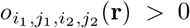 the ellipsoids should be updated such that the new value of 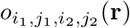 is 0. This is done by updating partly the positions and partly the shapes of the ellipsoids.

In practice, a small constant is added to *o* to make sure that there is a minimum distance between fibres in order to avoid overlaps in the final mesh-representation.

#### 2.2.4 Updating shapes and positions of fibre-ellipsoids

The key assumption, that axon morphology is influenced by the local environment [1][32], is implemented by requiring each axonal ellipsoid to adapt both shape and position according to its local environment in order to avoid overlapping structures. Each ellipsoid will have a specified deformation factor *δ*(∥**x**_*i,j*_(**r**) ∥) ∈ [0, 1] depending on the specified deformation factor map. Each fibre has a minimum allowed radius *μ*_*i*_ specifying that it should hold for any ellipsoid *j* that ∥**x**_*i,j*_(**r**) ≥∥ *μ*_*i*_. The overlap 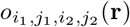 is divided in two equally big parts, and each part is handled by each of the two ellipsoids. So the deformation of an ellipsoid will result in a decrease in its size of at most 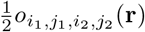. This will ensure that the ellipsoids will not have a negative overlap after being deformed. Since multiplying by a deformation matrix corresponds to a scaling along **r**, the distance that the extremal point should be moved has to be divided by the current size along **r** to get the scaling factor *s*. So in the case with ellipsoid *j*_1_ of fibre *i*_1_ the value of *s* would be 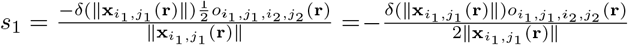 . In the case where this deformation would make the ellipsoid’s size smaller than the minimum *μ*_*i*_, a less negative value of *s* is chosen such that the resulting size is exactly 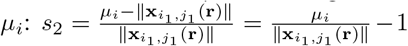. So the resulting value of *s* is *s* = max(*s*_1_, *s*_1_). So the deformation matrix **D**_1_ for ellipsoid *j*_1_ is **D**_1_ = max(*s*_1_, *s*_2_)**rr**^*T*^ + **I**. Now the shape matrix can be updated by calculating 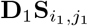 . The equivalent computations are performed for ellipsoid *j*_2_.

Secondly, the positions of the ellipsoids are updated. Each ellipsoid is treated as if they had equal ‘mass’, i.e. the centre of their positions is conserved. The total distance to move the ellipsoids apart is equal to the overlap 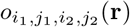after the shapes have been updated. By moving the two ellipsoids by the same amount, the new positions are updated to 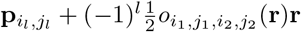 for *l* ∈ *{*1, 2*}*.

### 2.3 Post-processing

A phantom is outputted in two formats: an ellipsoid-representation and a mesh-representation. The ellipsoid-representation is the updated config-file where the shape and position of each ellipsoid is specified. This config-file can be edited and used as input for further optimisation. The mesh-representation is acquired from the ellipsoid-representation by generating tube-like mesh-segments to connect all ellipsoidal cross-sections within a fibre. The user specifies the number of radial segments that the tubes should have (radial resolution). The number of longitudinal segments is equal to the number of ellipsoids on the fibre. The cells are exported as they are i.e. meshes having the shape of the corresponding ellipsoids. The mesh-representation can be outputted as individual files for each fibre and cell, or as one combined file containing all elements.

#### 2.3.1 Optimisation of meshes for Monte Carlo diffusion simulations

The higher the ellipsoid-density, the smaller deviation is reached between the ellipsoid-representation and the mesh-representation due to gaps between ellipsoids. Hence, a high ellipsoid density is required to avoid overlapping fibres after conversion to the mesh-representation. This means that the resulting meshes have a likewise high longitudinal resolution. Especially if the meshes are intended for Monte Carlo simulations of dMRI, this is unfavourable since each mesh element comes with a computational cost due to the extensive collision-detection involved. Meanwhile, the high mesh resolution is often redundant.

By performing Garland Heckbert simplification [36] of each myelin and axon mesh, we reduce the number of faces and vertices significantly. The reduction depends on the initial resolution and the allowed deviation between the initial and the simplified mesh. We allow a deviation of `minDistance`*/*2 − 0.0005 *μ*m, where `minDistance` is the minimum distance allowed between ellipsoids of different fibres. For a radial resolution of 16 segments and `ellipsoidDensityScaler` of 0.20, the Garland Heckbert simplification decreases the number of faces by around 50%.

Before employing the phantoms in Monte Carlo diffusion simulations, the ends of the meshes are sealed to obtain “water tightness”.

The intra-axonal compartment can be extended by making two extra copies of the ellipsoids mirrored through each of the fibre’s endpoints without the requirement of further optimisation.

### 2.4 Morphological analysis

To demonstrate the biological relevance of the morphological features expressed in the numerical phantoms generated with the WMG, we compare the morphology of a set of numerical phantoms with real axons. The real axons originate from a monkey brain, and have been quantified by segmentation from XNH-volumes in previous work [1]. The meshes are available at www.drcmr.dk/powder-averaging-dataset. The morphological metrics quantify axon diameter variations, cross-sectional eccentricity, and tortuosity. All metrics are sampled with approximately 375 nm spacing along axon trajectories (some variation due to tortuosity).

#### 2.4.1 Axon diameters

We quantify axon diameter (AD) by the equivalent diameter as in [20]. I.e., axon diameters reported here are the diameter of a circle with an area equal to that of the axonal crosssection perpendicular to its local trajectory. For the real axons, the metric is extracted from the segmentation. This method is described in more detail in [1]. For the WMG-generated axons, the metric is extracted from the ellipsoid-representation.

For individual axons, mean AD (mean(AD)) is calculated over all measurements along an axon, while the standard deviation of the AD (std(AD)) quantifies the variation of these values.

#### 2.4.2 Cross-sectional eccentricity

We quantify cross-sectional eccentricity based on elliptic parameterisation by 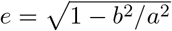 where *a* is the major axis and *b* is the minor axis (see Fig. 10). For the real axons, the circumference of the segmentation is sampled by 24 points. The length of the major and minor axes of a corresponding ellipse is then extracted by fitting a Principal Component Analysis to the sampled points. The results depend on the resolution and smoothing of the images. For the WMG-generated axons, the lengths of the major and minor axes are extracted directly from the ellipsoids.

For individual axons, mean eccentricity (mean(eccentricity)) is calculated over all measurements along an axon, while the standard deviation of the eccentricity (std(eccentricity)) quantifies the variation of these values.

#### 2.4.3 Tortuosity

We quantify tortuosity based on the tortuosity factor and the maximum deviation. The tortuosity factor is given as the fraction between the entire length of the centreline of an axon, and the length of a straight line going from end to end of the axon (see Fig. 11). The maximum deviation is given by the maximum distance between the centreline and the straight line (see Fig. 11).

### 2.5 Phantoms presented here

All phantoms included in this paper, are generated based on the optimisation scheme shown in Tab. 1. The growth in the first stage is rougher (i.e. lower `ellipsoidDensityScaler` and higher `growSpeed`), and then gets finer through the stages (i.e. higher `ellipsoidDensityScaler`, and lower `growSpeed`). Different optimisation schemes may yield different FVFs and generation times. The following parameters are kept constant and equal for all phantoms:

**Table 1:** Optimisation scheme used to generate the phantoms presented in the Results section.

| Stage | `ellipsoidDensityScaler` | `growSpeed` | `maxIterations` |
| --- | --- | --- | --- |
| 1 | 0.50 | 0.02 | 50 |
| 2 | 0.50 | 0.01 | 500 |
| 3 | 0.25 | 0.01 | 100 |
| 4 | 0.25 | 0.01 | 10 |

- `targetFVF = 0.8`
- `mapFromMaxDiameterToMinDiameter =`

`{‘from’ : [0.2, 0.5, 1.25], ‘to’ : [0.2, 0.2, 0.5]}`

Some phantoms include cells. The size and shape of the cell clusters are determined by randomly sampling the ellipsoid axes from normal distributions. Each primary axis *l*_1_ is sampled from the distribution *l*_1_ ∼ *N* (*μ* = 13 *μ*m, *σ* = 2 *μ*m), while the secondary *l*_2_ and tertiary *l*_3_ axes are set to be equal, and sampled from the distribution *l*_2_ = *l*_3_ ∼ *N* (*μ* = 5 *μ*m, *σ* = 1 *μ*m). This results in a mean fractional anisotropy of 0.54. The dispersion angles of the cell clusters (i.e. orientation relative to the primary axis of axons) are sampled from a uniform distribution over the interval (− 23, 23) deg. This is in accordance with observations of CVF, anisotropy and fibre dispersion in [1], where cell clusters were approximated as ellipsoids (or tensors).

## 3 Results

To demonstrate the applications of the interactive feature of the WMG, we focus on two scenarios: general cell mobility and inflammatory response. Firstly, we demonstrate the biological relevance of the morphological features expressed in the numerical phantoms generated with the WMG, by comparing the morphology of a set of numerical phantoms with real axons. The real axons originate from a monkey brain, and have been quantified by segmentation from XNH-volumes in previous work [1]. Then, by interactively changing the cell configurations over time, we show the morphology of the surrounding fibres changes consequently. We quantify the changes by analysing longitudinal axon diameter variations, longitudinal variations of cross-sectional eccentricity, and tortuosity of fibres.

### 3.1 The WMG mimicking different tissue compositions

The global fibre dispersion is controlled by the parameter *ϵ*. Examples of varying degrees of dispersion are shown both with and without cells in Fig. 4. When increasing *ϵ*, the fibres are forced to bend and deform more to achieve a higher FVF. Similarly, the presence of the static cells forces the fibres to adapt around them.

**Figure 4:**
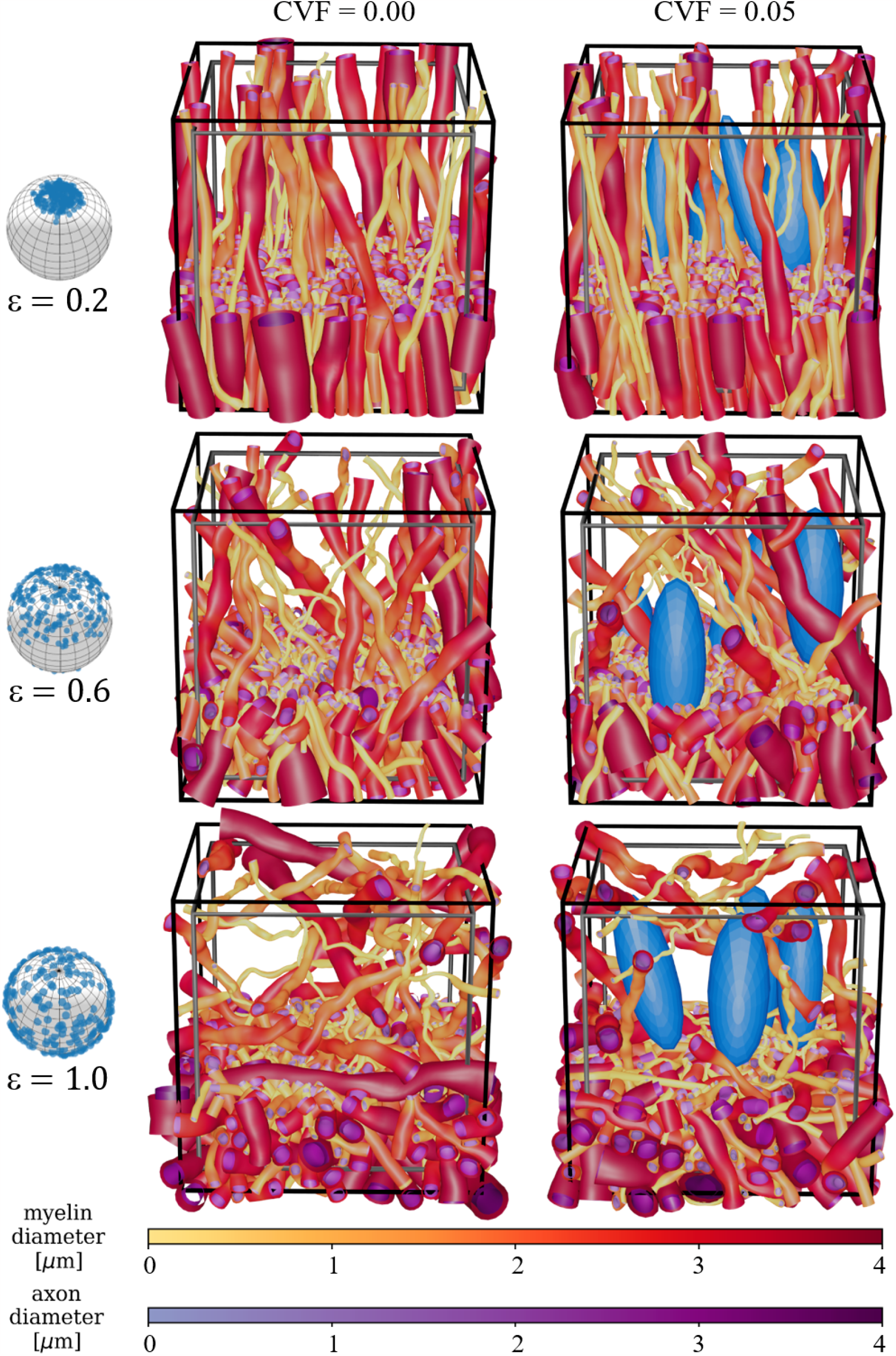
Demonstration of the global fibre dispersion parameter *ϵ* and the presence of cells (blue ellipsoids). The unit spheres on the left show the global dispersion associated with each epsilon. A cut is made at 1/3 of the voxel’s height to enhance the visualisation of individual fibre morphology. Above this height, the meshes are pruned such that only 7% are left. The black voxel marks the boundary of the ellipsoid centres, while the grey voxel marks the volume for which the FVF is optimised. The two voxels are used to avoid boundary effects within the optimised volume.

Crossing fibre bundles can be generated in two configurations sheets [33] and interwoven. Examples are shown in Fig. 5.

**Figure 5:**
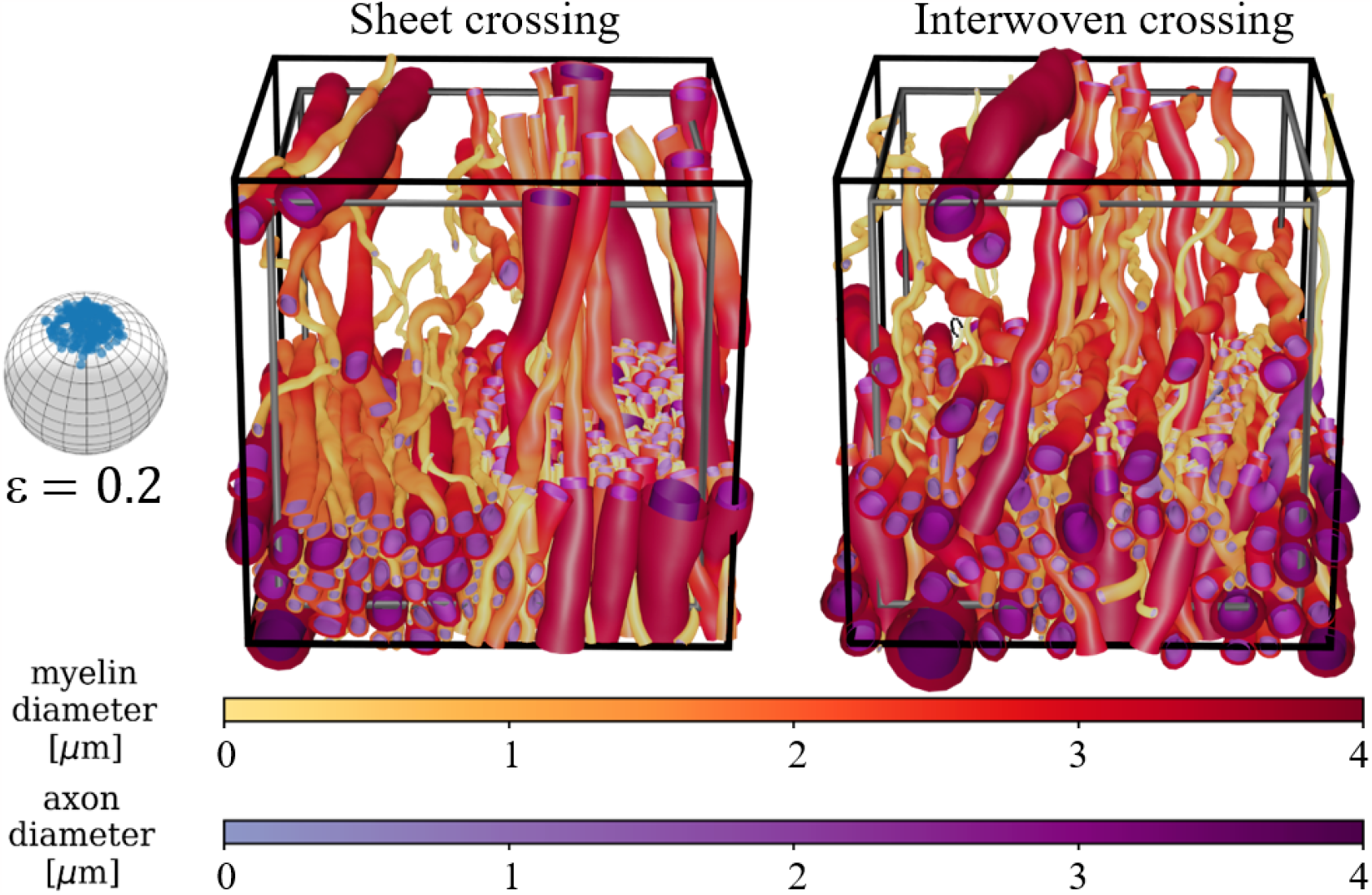
Demonstration of two crossing bundle configurations. The black voxel marks the boundary spatial limit of the ellipsoid centres, while the grey voxel marks the volume for which the FVF is optimised. In both examples, the number of fibres is split equally between the two perpendicular bundles. The split and fibre directions can be varied, and the number of crossing bundles can be increased. **a)** Sheet configuration where the fibres of each bundle are separated into sheets. **b)** Interwoven configuration where the fibres of each bundle are interwoven with each other.

Demyelination can be mimicked by selectively stripping the myelin from axons and adding cells to mimic an inflammatory response as shown in Fig. 6. This can be done either between optimisation steps by decreasing the allowed diameter range of a fibre to that of the corresponding axon, or by removing the myelin mesh after optimisation.

**Figure 6:**
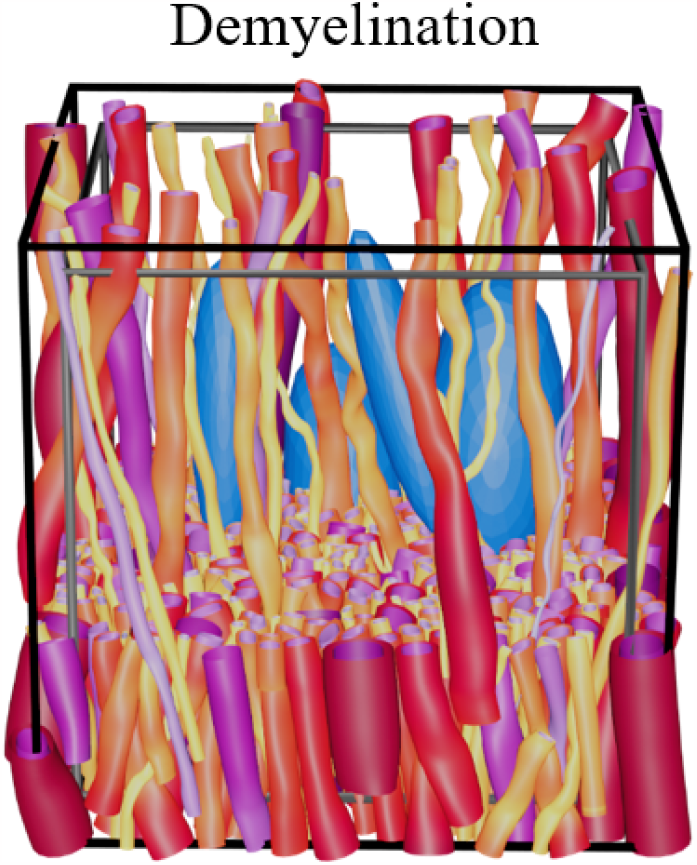
Demonstration of demyelination showing a demyelinated version of the phantom shown in Fig. 4 (*ϵ* = 0.2, with cells). The black voxel marks the boundary for the ellipsoid centres, while the grey voxel marks the volume for which the FVF is optimised.

### 3.2 Performance of the WMG

Fig. 7 shows the obtained FVFs and the processing time for the parameters and optimisation scheme described in Tab. 1. We achieve mean(FVF) ≥ 0.72 for phantoms of all degrees of *ϵ* without cells and mean(FVF) ≥ 0.66 with cell clusters (CVF=0.05). It is seen that the processing time increases with *ϵ*. This is a consequence of the increased fibre dispersion leading to a higher frequency of collisions of the chain of ellipsoids comprising the axons. Each collision necessitates correction, leading to a longer optimisation time. Likewise, the increased dispersion complicates dense packing and results in lower FVF. Similarly, the presence of cells (CVF=0.05) results in a much increased processing time compared to phantoms which have otherwise identical parameter configurations. This is due to the additional volume taken up by the static cell clusters further complicating the packing and resulting in lower FVF.

**Figure 7:**
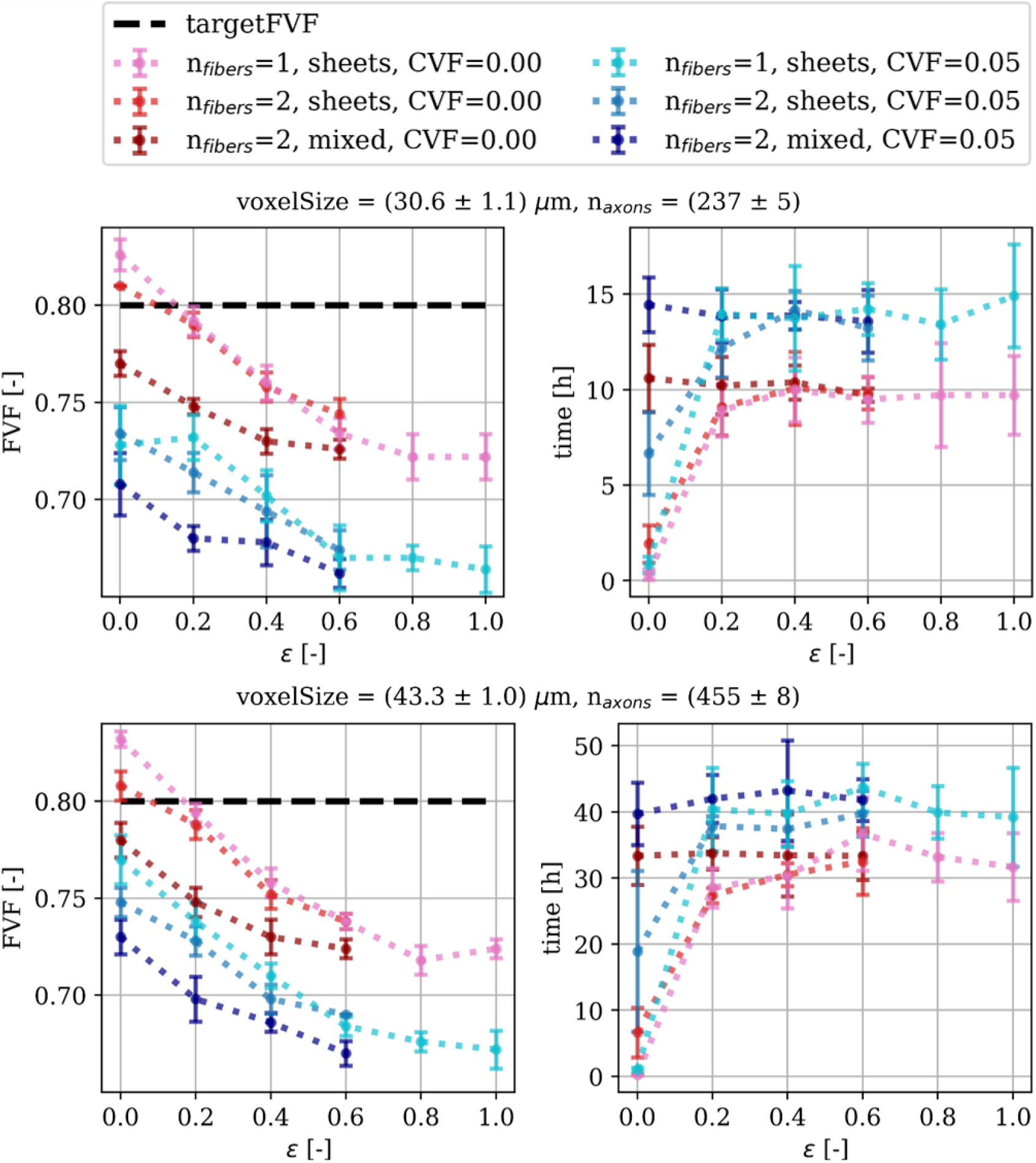
Performance of the WMG in terms of achieved axonal volume fraction (FVF) and processing time. Each point is based on 5 samples/repetitions. mean(FVF)>0.66 is achieved for all configurations. The complexity of the phantom configuration is increased by increasing dispersion (*ϵ*), adding cell clusters, and introducing fibre crossings. The higher complexity makes it more challenging to achieve higher FVFs.

### 3.3 Morphological variation of WMG-generated phantoms

To demonstrate that the morphological features expressed in the WMG-generated phantoms are relevant in relation to real 3D morphology, we compare the morphology of the axons with real axons quantified from XNH-volumes. The comparisons are based on longitudinal axon diameter variations, cross-sectional eccentricity, and tortuosity all metrics which are reflected in the dMRI signal.

#### 3.3.1 Longitudinal axon diameters

Fig. 8 shows an example of target and output distributions of mean axon diameters for individual axons. Much flexibility in morphology parameters is required to obtain the high FVFs. Hence, some deviation from the target is to be expected. For the parameters and optimisation scheme described in Tab. 1, the mean *μ* values of the distributions are within 0.6 *μ*m of that of the desired distribution with *μ* = 1.8*μ*m.

**Figure 8:**
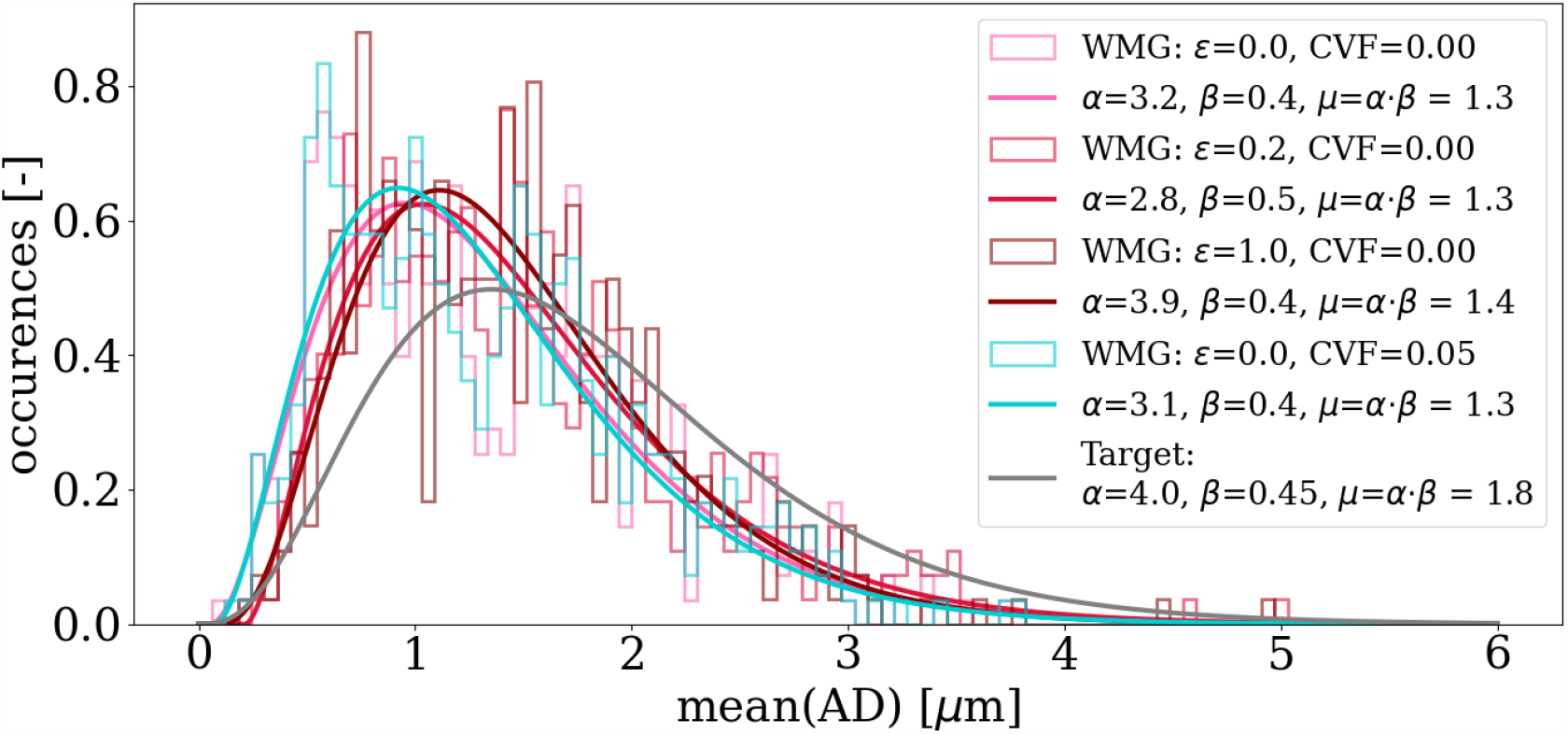
Target vs. outputted distributions of mean axon diameters for individual axons *mean*(*AD*). Gamma distributions are fitted to the histogram bins of the outputted mean axon diameters. The desired diameter of each axon is sampled from a gamma distribution (the desired distribution). Due to the packing procedure, some deviation is expected for the outputted distributions. Here, deviations of mean values are within 0.6 *μ*m.

Fig. 9 shows a positive correlation between the std(AD) and mean(AD). We compare with the XNH-samples and see an overlap of the metrics and a likewise positive correlation. We see that std(AD) can be regulated both by varying *ϵ* and CVF. The higher *ϵ* forces the individual axons to deform more to pack to the dispersing environment - and vice versa. Similarly for increased CVF (see “*ϵ* = 0.0, CVF=0.00” vs. “*ϵ* = 0.0, CVF=0.05”), where the presence of cells force additional ellipsoid deformation in order to obtain the dense packing.

**Figure 9:**
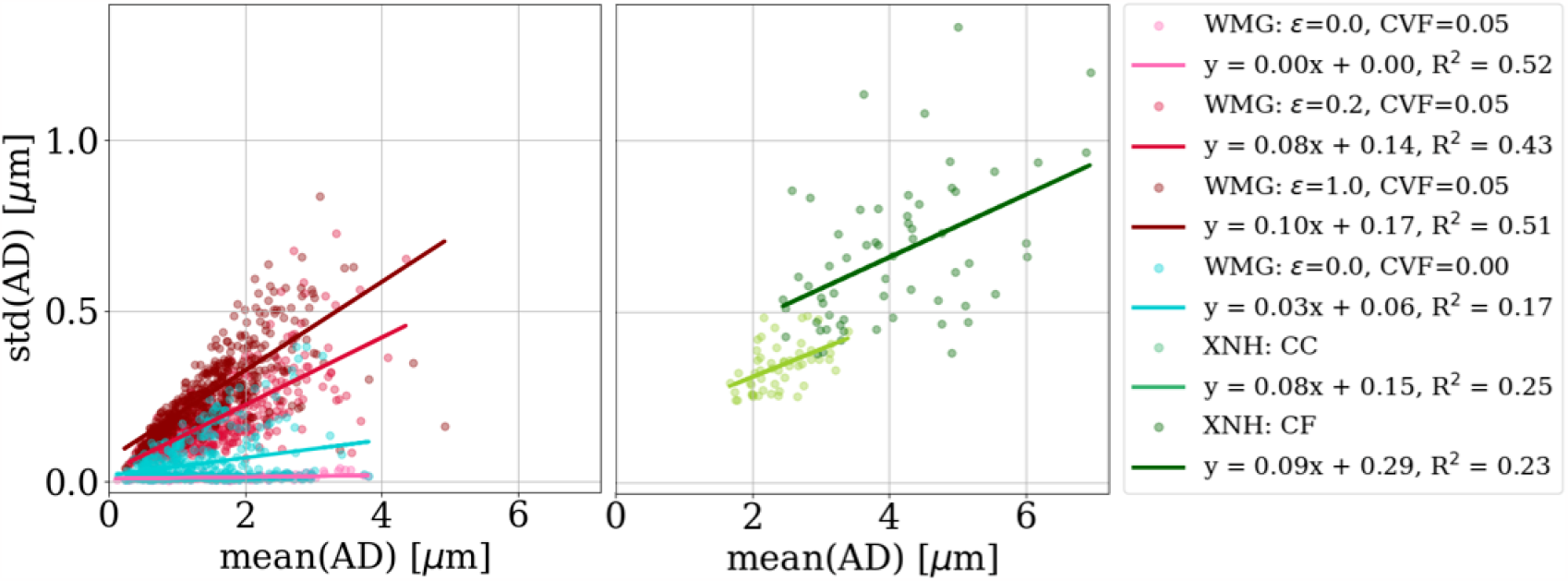
Longitudinal axon diameter variation. Each marker represents one axon from a given sample, while the lines are the linear fits of all axons in a sample/numerical phantom. The standard deviation of the mean axon diameter per axon (std(AD)) correlates positively with the mean axon diameter per axon (mean(AD)) - both for the XNH and WMG axons. Note that the distribution of axon diameters of the XNH axons is not representative of the tissue as a whole, but biased towards the larger axons due to those axons being more clear on the XNH images.

#### 3.3.2 Axonal cross-sectional eccentricity

Fig. 10 shows the variation in cross-sectional eccentricity along WMG-generated axons as a function of the mean cross-sectional eccentricity along the axons. Both the mean(eccentricity) and the std(eccentricity) can be regulated by varying *ϵ* and CVF. Again, because the higher *ϵ* forces the individual axons to deform more to pack to the dispersing environment – and vice versa. Similarly for increased CVF (see “*ϵ* = 0.0, CVF=0.00” vs. “*ϵ* = 0.0, CVF=0.05”), where the presence of cells force additional ellipsoid deformation to obtain the dense packing. The types of eccentricities in the XNH-segmented axons and the WMG-generated axons are different in nature. While the cross sections of the XNH axons are asymmetric and squiggly by nature, the cross sections of the synthetic WMG axons are limited to being elliptic due to the axon being comprised of ellipsoids. Hence, while the eccentricity measure does provide a rough estimation of the dominating cross-sectional eccentricity in both cases, it is not directly comparable across the two.

**Figure 10:**
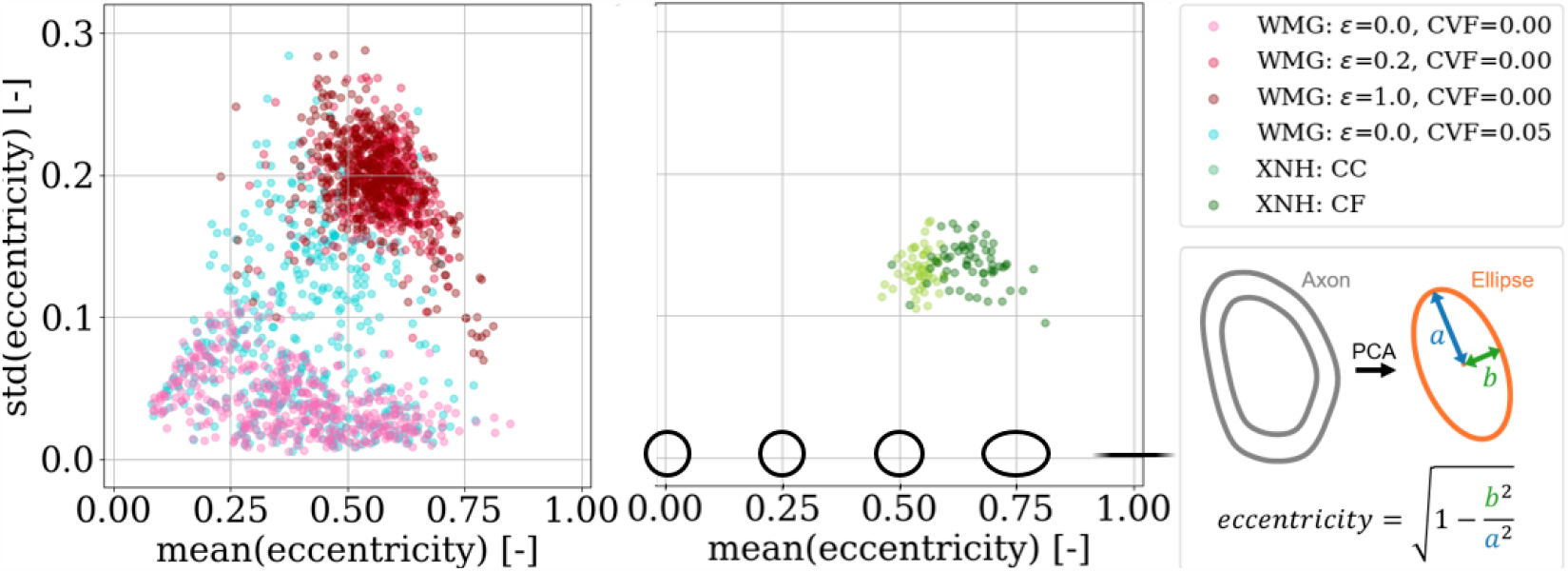
Cross-sectional eccentricity. We achieve an overlapping degree of eccentricity for the WMG-generated axons as we observed for the XNH axons. Meanwhile, the std(eccentricity) shows a higher degree of variation than the XNH samples. We see a higher degree of eccentricity in the crossing fibre (CF) compared to the corpus callosum (CC) quantified from the XNH images.

**Figure 11:**
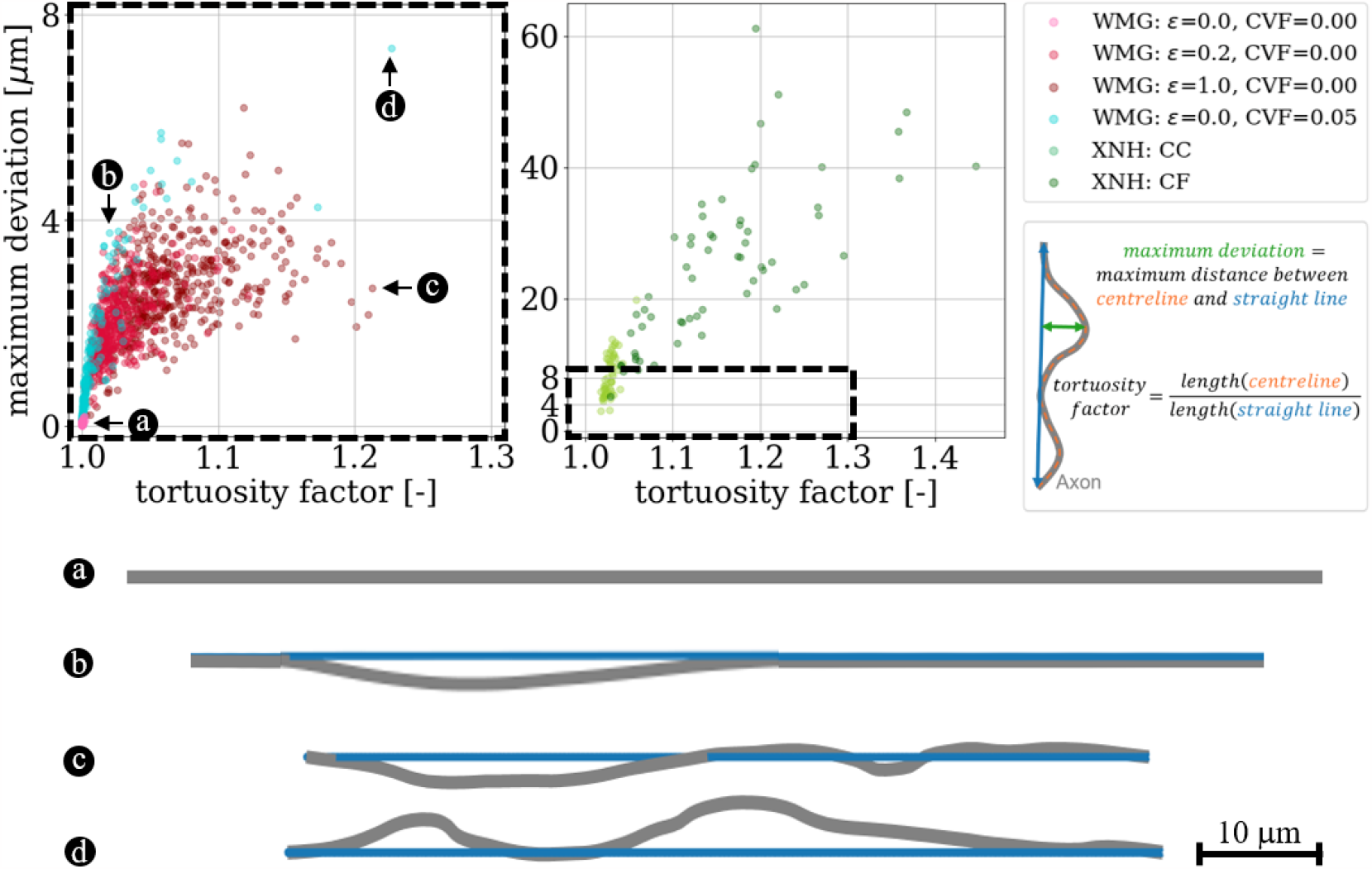
Tortuosity. We quantify the tortuosity of the axons based on the tortuosity factor and the maximum deviation (both metrics are explained in the figure) to compare with the real axons segmented from XNH-imaging of corpus callosum (CC) and crossing fibre (CF) from a monkey. It is possible to reach overlapping tortuosity factors for the synthetic WMG axons as is observed for the real XNH axons from the corpus callosum (CC) region. However, we do not match the high tortuosity factors and maximum deviations observed for the crossing fibre (CF) region. a, b, c, and d show examples of 2D projections of the axonal centrelines at different ends of the spectrum.

#### 3.3.3 Axonal tortuosity

A positive correlation between maximum deviation and tortuosity both for axons from the WMG and XNH is observed. This is shown in Fig. 11. The tortuosity can be regulated by the dispersion parameter *ϵ* and by adding cells. The higher dispersion and CVF mean that the axons have to bend more around each other to not overlap and vice versa.

### 3.4 Interactive configuration allows the mimicking of dynamic environments

Dynamic cells are mimicked by interactively changing the cell configuration during phantom optimisation and letting the fibres adapt accordingly. Fig. 12 shows how a dynamic cell environment can influence the morphological metrics of the surrounding axons. In the right column, we see how moving cells are causing an increase in axon diameter variations, eccentricity, and tortuosity over time. In Fig. 13, we see a similar trend when the CVF is increased over time except that in this example, the axons are generally straight and have non-eccentric cross-sections at the first time point due to CVF=0.00.

**Figure 12:**
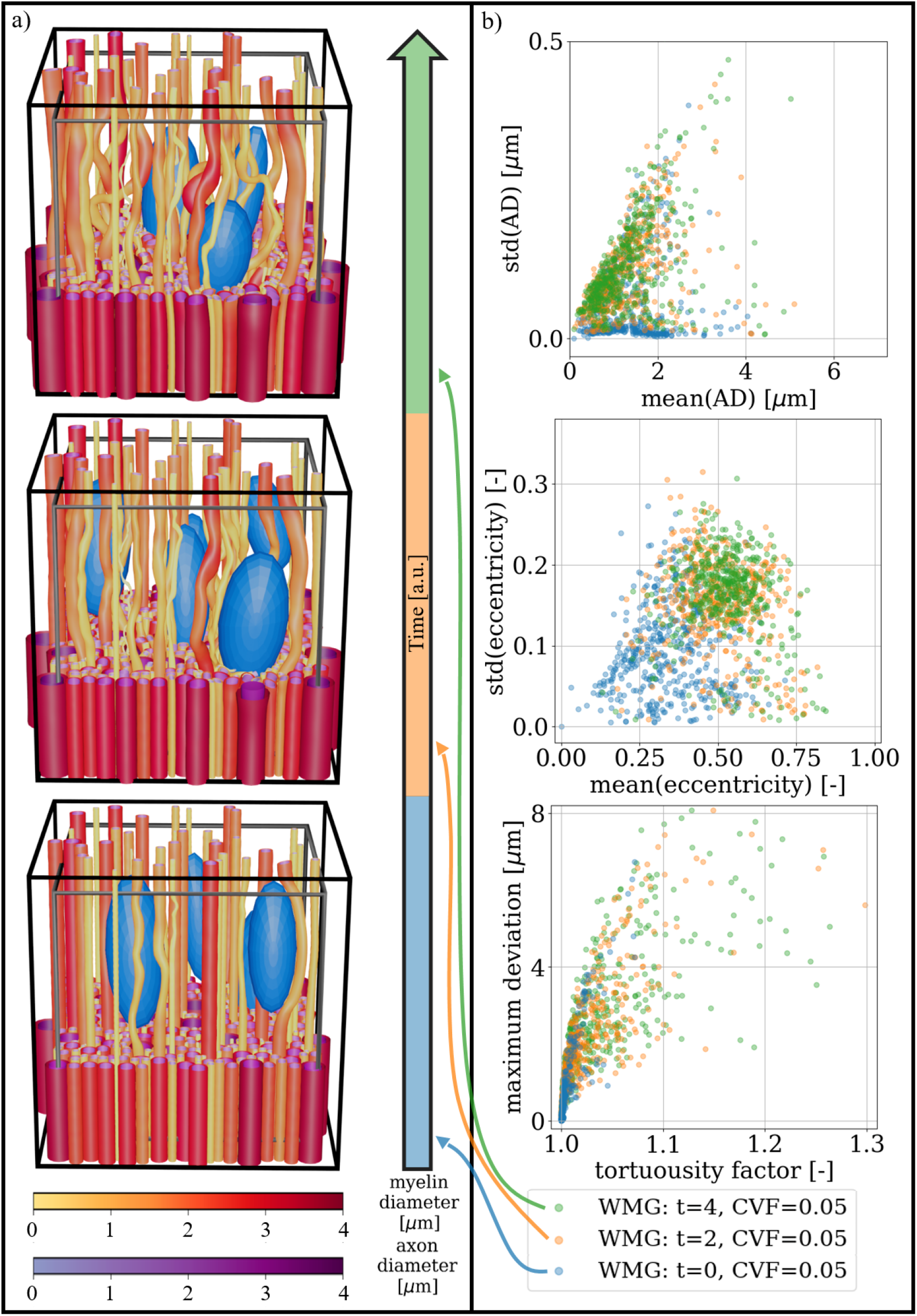
General cell mobility: The CVF is kept constant while cells are moved around over time. **a)** Visualisation of three temporal snapshots of a phantom. A cut is made at 1/3 of the voxels height to enhance the visualisation of individual axon morphology. Above this height, the meshes are pruned such that only 7% are left. The black voxel marks the boundary of the ellipsoid centres, while the grey voxel marks the volume for which the FVF is optimised. **b)** Morphological metrics computed for the phantom snapshots seen in a). When starting from otherwise straight axons, the moving cells cause increasing diameter variations (top), eccentricity (middle), and tortuosity (bottom) over time. For an explanation of metrics for eccentricity and tortuosity, see Fig. 10 and Fig. 11, respectively.

**Figure 13:**
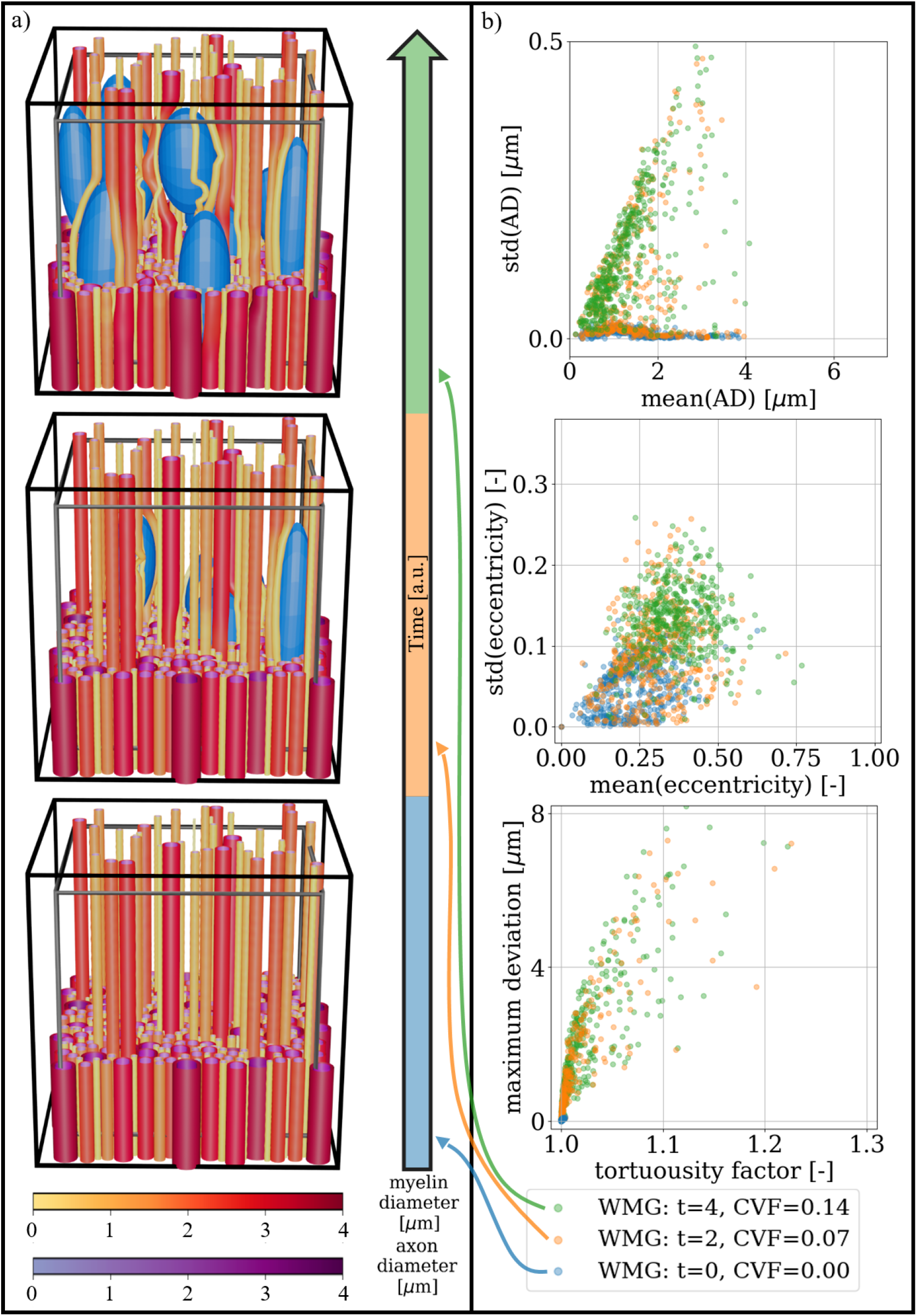
Inflammatory response: The CVF is increased in steps of 0.035 from 0.00 to 0.14 over time. **a)** Visualisation of three temporal snapshots of a phantom. A cut is made at 1/3 of the voxels height to enhance the visualisation of individual axon morphology. Above this height, the meshes are pruned such that only 7% are left. The black voxel marks the boundary of the ellipsoid centres, while the grey voxel marks the volume for which the FVF is optimised. **b)** Morphological metrics computed for the phantom snapshots seen in a). When starting from otherwise straight axons, the increasing cell fraction causes increasing diameter variations (top), eccentricity (middle), and tortuosity (bottom) over time. For an explanation of metrics for eccentricity and tortuosity, see Fig. 10 and Fig. 11, respectively.

## 4 Discussion

With the development of the WMG tool, we enable the generation of interactive numerical white matter phantoms composed of the compartments of axons, myelin, cells and extracellular space. This allows us to mimic the dynamic nature of white matter by interactively changing the configuration of the cell clusters within a given phantom while the surrounding axon morphology adapts accordingly. With these phantoms, we can thereby analyse how specific cell dynamics (such as mobility and inflammation [3][4][5]) influence the morphology of the surrounding axons. Furthermore, the very same phantoms can be employed as mesh inputs for Monte Carlo diffusion simulations. This enables us to study how these finer detailed morphological changes affect the dMRI signal and to evaluate microstructure models.

When comparing morphological metrics between axons generated with the WMG tool and real axons segmented from XNH-imaging of a vervet monkey brain (corpus callosum and crossing fibre), we see an overlap with the metrics of axon morphologies from the corpus callosum. However, the WMG tool does not offer the degree of tortuosity observed for the crossing fibre region.

Based on the assumption that real white matter morphology carries a history of dynamic events, the WMG tool provides an improved understanding of the dynamics of white matter morphology - both for healthy and diseased tissue.

### 4.1 WMG-generated axons possess histology-resembling morphological metrics

We analysed the morphology of the individual axons from phantoms by the WMG tool. Based on the metrics for longitudinal diameter variations (Fig. 9), longitudinal variations of cross-sectional eccentricity (Fig. 10), and tortuosity (Fig. 11), we show that the WMG tool is capable of generating axons with morphology closely resembling that observed for axons from the corpus callosum of a vervet monkey brain with XNH imaging in [1][32]. These morphological metrics have been found to influence the dMRI signal from individual axons [16][17][1][32][18][21], and are hence essential when generating numerical white matter phantoms with the purpose of evaluating microstructure models for dMRI.

The longitudinal axon diameter variation (Fig. 9, y-axis) and the longitudinal variations of cross-sectional eccentricity (Fig. 10), and the tortuosity factor (Fig. 11, x-axis) reached by the WMG tool, are in agreement with that of the XNH-imaged corpus callosum. The maximum deviations (Fig. 11, y-axis), however, are generally much higher in the XNH-imaged corpus callosum. Meanwhile, the XNH-imaged crossing fibre is more challenging to match. Here, only the degree of eccentricity (Fig. 10) is in agreement. To obtain a higher degree of tortuosity, and especially the maximum deviation, the ellipsoid chains that make up the foundation of the axons would have to be initialised with an initial tortuosity rather than in a straight line. While this is not a standard configuration of the WMG tool, it can be obtained through manual configuration. No other tools for the numerical synthesis of white matter phantoms are known to output this degree of tortuosity either.

Because the low signal-to-noise ratio of the XNH volumes challenged a robust manual segmentation of axons with mean diameters smaller than 2 *μ*m, the segmented axons are not representative of the underlying distribution w.r.t. diameter [1]. This quantification does, however, provide valuable insight into the realistic range of morphological metrics for the larger axons.

For extra-cellular space, we targeted a volume fraction of 0.2 (i.e., targetFVF=0.8) in agreement with what has been reported for normal adult brain tissue - which is between 0.15 and 0.30, and typically 0.20 [37]. Fig. 7 shows that all WMG phantoms without cells, reach well within the targeted range for all degrees of fibre dispersion. More precisely, we obtain FVFs between 0.83 *±*0.00 and 0.80 *±*0.01 for single fibre bundle phantoms with dispersion between 0 deg and 18 deg (*ϵ* of 0.0 and 0.2, respectively), and FVF of 0.72 *±*0.01 for dispersion 90 deg (*ϵ* of 1.0). Meanwhile, the MEDUSA tool [28] reaches FVFs between 0.72 and 0.62 for single-fibre-bundle phantoms with dispersion between 0 deg and 20 deg, and the ConFiG tool [29] reaches FVFs between 0.75 and 0.71 for single-fibre-bundle phantoms with dispersion between 7.75 *±*4.23 deg and 17.46*±* 9.92 deg. Although the CACTUS tool [30] reaches impressive FVFs between 0.91 and 0.95 for single-fibre-bundle phantoms with dispersion between 0 deg and 25 deg, such high volume fractions are not necessarily relevant.

### 4.2 Axon morphology is modulated by cell dynamics

By exploiting the interactive component of the WMG tool to mimic cell dynamics in the form of general cell mobility and inflammatory response, we show how axon morphology can be modulated over time by these expected dynamics (Fig. 12 and Fig. 13). Under the assumption that real white matter follows similar principles, we can thereby use the WMG tool to study expected tissue response in various physiological or pathological scenarios involving dynamic cell behaviour. No other tool offers this feature.

Cell dynamics are continuously occurring in healthy tissue in the form of glial cell mobility [3][4][5]. However, because these changes are homogeneously distributed across the tissue, their influence on morphological metrics should stabilise over time and not cause net changes to the dMRI signal. Here, we mimic cell mobility by changing the configuration of cells over time while keeping the CVF constant (Fig. 12). By starting from straight axons, we see how the continuous cell mobility induces an increase in longitudinal morphological variation as the axons adapt to the changing environment.

### 4.3 Synthesis method and tissue configurations

The high morphological flexibility enabled by the ellipsoidal building blocks of the WMG tool allows for a large variety of different tissue configurations. Here, we show examples of varying dispersion (Fig. 4), fibre crossings (Fig. 5), demyelination (Fig. 6), inclusion of static cells (Fig. 4), cell mobility (Fig. 12), and inflammation (Fig. 13). A key assumption for the WMG tool is that axon morphology is modulated to the local environment, and thereby adapting according to surrounding axons and cells.

The WMG tool is the first tool developed for generating numerical white matter phantoms which includes an interactive component to allow the mimicking of cell dynamics. However, the tools MEDUSA [28], ConFiG [29], and CACTUS [30] do excel when it comes to computational efficiency and more complex morphological features.

Our concept of ellipsoidal building blocks is very similar to that of MEDUSA [28] where spherical building blocks are used. While the sphere representation allows for a high representational power of the longitudinal axon morphology characteristics, it does not allow for eccentric cross-sections as documented by histology [1][19][20]. On the other hand, the ellipsoid geometry allows for more degrees of freedom which, in turn, enables eccentric cross-sections although, still lacking cross-sectional squigglyness to truly resemble histology. Both tools are based on variations of a force-biased packing algorithm, first introduced by [34], where the phantoms are obtained as an equilibrium between repulsion forces (for avoiding overlapping axons and cells) and recover forces (for ensuring the structure of the individual axons). Similar mechanics are used to shape the axons in CACTUS [30], where each axon is represented by capsular building blocks during the optimisation of axon trajectories. However, the CACTUS tool contains an additional step of radial optimisation, which can increase packing densities and increase the realism of the axonal cross-sections by introducing the desired natural squigglyness. Meanwhile, the ConFiG tool [29] focuses on the initial growth and maturation of white matter by mimicking how axons are guided by chemical cues and adapt to the available space as they grow and extend from one end to the other. Hence, rather than initialising all axons along straight lines that extend the entire length of the voxel as done in the other tools, each axon is grown stepwise from one end to the other. This is followed by a radial optimisation step similar to that of CACTUS [30]. A similar finishing radial optimisation could beneficially be applied in the WMG tool to obtain likewise more realistic cross-sections.

### 4.4 Limitations and future work

#### 4.4.1 Tissue representativity

Numerical phantoms from the WMG tool can include axons, myelin, and cell clusters. These are the most prominent structures in brain white matter. However, additional structures present in the brain white matter might affect water diffusion and hence the dMRI signal. Such structures include microtubules, mitochondria, nodes of Ranvier, neuron cell bodies, and dendrites. These will be implemented in a future version of the WMG tool.

With the WMG tool, the g-ratio is modelled as constant throughout the axon length. However, it has been found that this ratio can vary across myelin segments [1].

Demyelination in the WMG tool is mimicked by randomly stripping myelinated axons from their myelin sheets and by increasing CVF. The intermediate steps of the demyelination process are not modeled.

The size of the phantoms is crucial for the representational power of tissue features. For the WMG tool, the processing time is the limiting factor. It has been shown that phantom sizes larger than (200 *μ*m)^3^ can reduce the sampling bias [15] and hence improve the representational power. Moreover, while efforts are made to minimize structural boundary effects during synthesis, complete avoidance is challenging. Furthermore, when conducting Monte Carlo diffusion experiments, it is necessary to initialise the particles with an additional distance from the boundary to prevent probing the ends of the axons. For ex vivo diffusivities (0.6 *μ*m^2^ms^−1^), a 20 *μ*m boundary has been applied to avoid this [1][32][21]. For in vivo diffusivities (2 *μ*m^2^ms^−1^), a 30 *μ*m boundary has been applied [32].

The WMG tool does offer a way to extend the intra-axonal compartment. All axon and myelin meshes can be mirrored around voxel boundaries without the requirement of further optimisation. However, the added amounts of meshing do come with a computational cost when applied in Monte Carlo diffusion simulations by threefold.

#### 4.4.2 Improving the computational efficiency

Creating phantoms is currently a time-consuming process with 28 *±* 3 hours spent on generating phantoms containing a single fibre direction with *ϵ* = 0.2 and 55*±* 8 axons in a voxel size of 43.3 *±* 1.0 *μ*m using one core (Fig. 7). In comparison, for generating a phantom of comparable configuration, the parallelisable CACTUS tool [30] requires 4 hours for generating a phantom of 33478 axons in a voxel size of (500 *μ*m)^3^ using 64 cores. However, since the optimisation procedure for the WMG tool relies on checking overlaps between ellipsoids, and this is formulated with linear algebra, there presents an excellent opportunity for leveraging GPUs to enhance efficiency. Therefore, future efforts will focus on implementing the WMG tool in a GPU-compatible manner to accelerate the phantom generation.

While it is already possible to generate large and numerous phantoms with the WMG tool, addressing the computational efficiency will significantly expand the capacity. This enhancement will be crucial for facilitating the generation of large-scale and machine learning-friendly datasets.

#### 4.4.3 Mimicking compression and stress of fibrous tissues

The interactive config-files and the force-biased packing algorithm of the WMG tool could further be applied to mimic the compression and stress of tissue. Such events can be mimicked by applying selective stretch or compression to a voxel and its content through interactive changes to the config-file. Meanwhile, cell elements can be added to mimic the accompanying necrotic debris and cellular immune response. Applications lie within traumatic brain injuries where tissue compression often arises due to external forces, and/or swelling in one area causing compression in adjacent regions [38]. But the applications also expand to dynamic responses in other types of fibrous tissues. Tissues like muscle, tendon, and ligament are prone to traumatic pulls causing damage and even tearing of their fibres.

Proper realignment of damaged fibres is crucial for regaining strength and function [39]. A quantitative evaluation of compression and stress of fibrous tissues could thereby make a valuable contribution to this broader research community.

## 5. Acknowledgements

S.W. has been supported by a DTU Alliance Stipend [project number 102682]. S.W., M.A., and T.B.D. have received funding from the European Research Council (ERC) under the European Unions Horizon Europe research and innovation programme (grant agreement No. 101044180) (Principal Investigator: T.B.D.). O.P., M.A., and H.M.K. have been supported by Capital Region Research Foundation Grant [grant number A5657; principal investigator: T.B.D.].

## 6. Contributions

O.P., M.A., J.A.B., and T.B.D. developed the concept. O.P. implemented the framework and the graphical user interface in TypeScript. S.W. implemented configurations for the different use cases, morphological analysis, and bridging to Monte Carlo diffusion simulations in Python 3. H.M.K. contributed to the morphological analysis. O.P., J.A.B., and S.W. wrote the methods section. S.W. wrote the remaining paper and created the figures. T.B.D. supervised the project. O.P., S.W., M.A., H.M.K., J.A.B., and T.B.D. revised the paper.

## 7 Data availability

The framework will be made available at https://white-matter-generator.github.io/. The code to reproduce the configurations and morphological analyses presented here will be made available at GitHub upon publication. The datasets generated and analysed during this study will be made available in the “White Matter Generator” repository at www.drcmr.dk/map-datasets upon publication.

## Notes

### Competing Interest Statement

The authors have declared no competing interest.

